# Immortalized AT2 Cell Lines from Healthy and IPF Lungs Enable 2D and 3D Cultures

**DOI:** 10.64898/2026.03.09.709900

**Authors:** Anas Rabata, Yujie Qiao, Xuexi Zhang, Jiurong Liang, Dianhua Jiang

## Abstract

Idiopathic pulmonary fibrosis (IPF) is characterized by impaired alveolar type 2 cell regeneration. However, robust in vitro models of human distal lung epithelium are limited. In this study, we generated immortalized AT2 cell lines from healthy and IPF lungs using HTII-280 sorting and SV40 large T antigen transduction. These lines retain key features of alveolar epithelial biology in both 2D and 3D cultures, including self-renewal, differentiation, and transitional cell states. They form 3D organoids efficiently under optimized feeder-free, serum-free medium conditions, with higher colony-forming capacity in healthy AT2 cell lines comparing with IPF AT2 cell lines. These accessible models recapitulate alveolar epithelial biology, offering a platform for cell-biology research and therapeutic development in lung diseases.

## To the Editor

Idiopathic pulmonary fibrosis (IPF) is a fatal interstitial lung disease characterized by repeated alveolar epithelial injury, defective repair, progressive scarring, and loss of lung function (*1*). Therapeutic options remain limited because the underlying mechanisms are poorly understood. Alveolar type 2 (AT2 or AEC2) cells, which produce pulmonary surfactant, are recognized as essential stem cells that maintain and repair the distal lung epithelium through self-renewal and differentiation (*2*). We and others have shown that AT2 regeneration fails and AT2 cell numbers are reduced in IPF, contributing to disease progression (*3*). Therefore, establishing robust in vitro models that accurately recapitulate these diseases is essential for uncovering mechanisms of distal lung pathology and enabling the development of new therapeutic strategies.

The isolation of primary AT2 cells has been well developed by many groups (*4-6*). Primary AT2 cells are notoriously difficult to maintain in culture. Although immortalized human airway epithelial cell lines have been widely used for in vitro lung research due to their availability, scalability, and robust two-dimensional (2D) and 3D growth, comparable models from the distal alveolar epithelium remain scarce. This limitation reflects persistent challenges, including the low yield of primary alveolar cells, poor adherence and instability in 2D monolayers, dependence on complex co-culture systems for 3D organoids, and difficulty preserving alveolar-specific identity over time.

The goals of this study are: (1) to establish immortalized AT2 cell lines from healthy donors and IPF explants, (2) to establish 2D monolayer cultures, and (3) to construct culture conditions to maintain 3D organoid cultures.

In this study, we successfully established immortalized AT2 cell lines from both healthy and IPF lung tissues. EpCAM^+^ epithelial cells were isolated from both healthy and IPF lung samples via enzymatic digestion and magnetic bead-based enrichment. Cells were initially cultured under conditions containing Rho-associated kinase (ROCK) inhibitor, Y-27632 (*7, 8*), to promote attachment and survival. These cultures were then transduced with lentivirus expressing SV40 large T antigen (SV40 LgT) for immortalization (*9*). To further purify AT2 populations, immortalized EpCAM^+^ cells were sorted by FACS using the specific AT2 membrane marker HTII-280, and the sorted cells were re-transduced with SV40 LgT to establish stable cell lines (Figure 1A and B). We noted that cells transduced only once with SV40 large T antigen (SV40 LgT) began to adopt a more spindle-like morphology and lose their triangular shape at higher passages. In contrast, cell lines subjected to a two-step SV40 LgT transduction process remained stable, retaining their morphology even at higher passages. Cell lines derived from IPF lungs required more time to reach confluence and showed reduced attachment to the culture plate; however, they exhibited similar features and morphology in 2D culture (Figure 1C).

**Figure 1:**
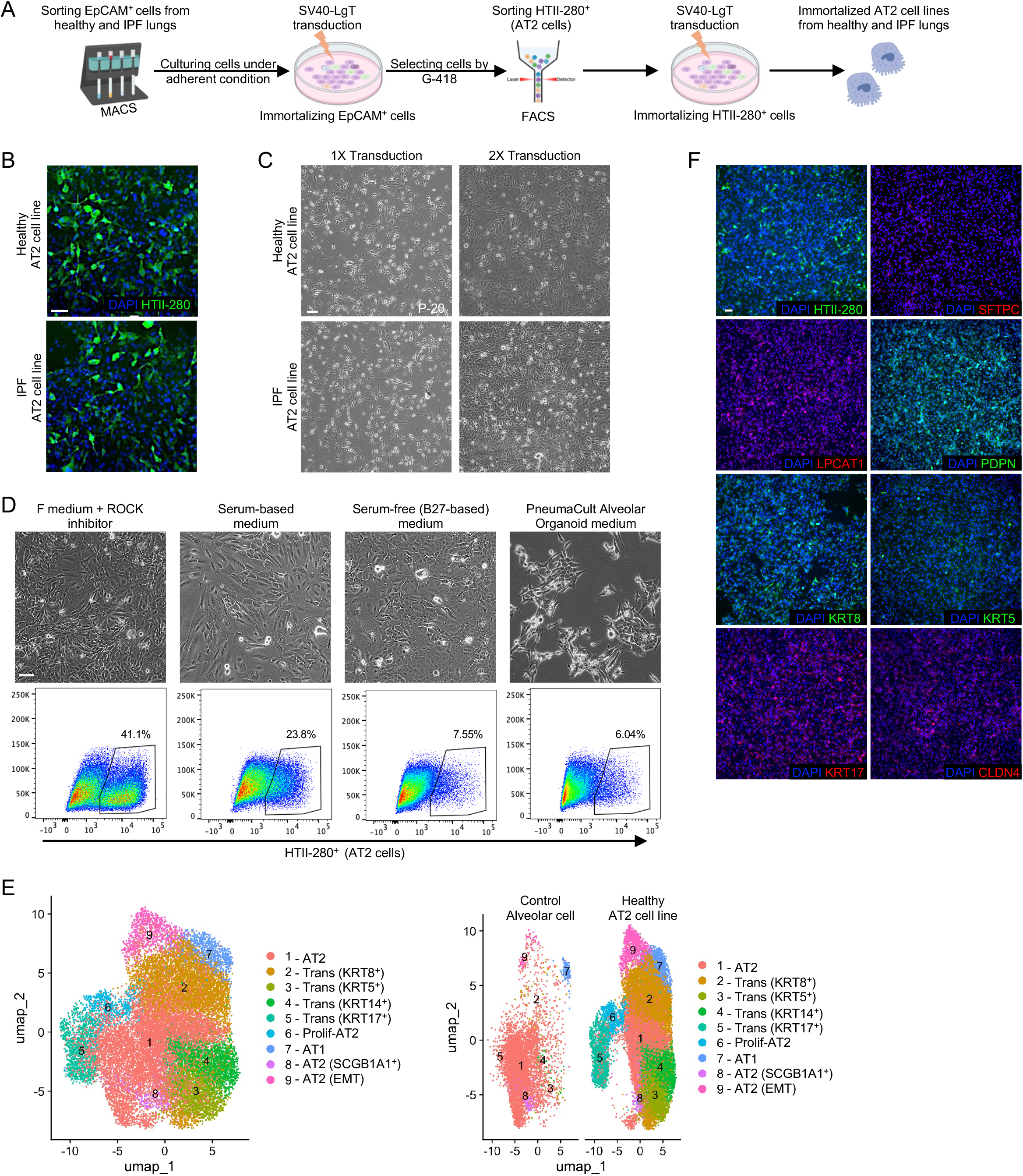
Generation of immortalized AT2 cell lines. (A) Experimental layout for generating immortalized AT2 cell lines from healthy and IPF lungs. (B) Immunofluorescence staining in 2D cultures showing expression of AT2 cell marker HTII-280 (green) in sorted cells from healthy and IPF immortalized AT2 lines. Scale bars, 100 μm. (C) Representative images of healthy and IPF AT2 cell lines at passage 20, following one or two rounds of SV40 large T antigen (SV40 LgT) transduction. Scale bars, 100 μm. (D) Representative images and Flow-cytometry gating strategies for immortalized HTII-280^+^ AT2 cells sorted from healthy EpCAM^+^ cells and cultured in 2D under deferent medium conditions: 1- F medium + ROCK inhibitor (3:1 DMEM/F12 to DMEM, 5% FBS, 0.4 μg/mL hydrocortisone, 5 μg/mL insulin, 8.4 ng/mL cholera toxin, 10 ng/mL EGF, and 10 μM Y-27632), 2- Serum-based condition medium (DMEM/F12, 10%FBS, 1X ITS and 10 μM Y-27632), 3- Serum-free (B27-based) medium (DMEM/F12 supplemented with B27 and 10 ng/mL EGF + ROCK inhibitor), 4- PneumaCult Alveolar Organoid medium. Scale bars, 100 μm. (E) UMAP plots of flow-enriched HTII-280^+^ cells from healthy AT2 cell lines and the data integrated with published primary AT2 cell data (GSE132915) as a control, showing clusters corresponding to AT2-like, AT1-like, and transitional cells. (F) Immunofluorescence staining of Healthy AT2 cell lines in 2D culture, showing expression of AT2 cell markers (HTII-280, SFTPC, and LPCAT1), AT1 cell marker (PDPN), and transition cell markers (KRT8, KRT5, KRT17, and CLDN4). Scale bars, 100 μm.

We compared the culture medium conditions (F medium + ROCK inhibitor) described by the Zea Borok laboratory (*8*) with three other formulations: a serum-based medium (*3*), DMEM/F12 supplemented with B27 and EGF + ROCK inhibitor (*10, 11*), and PneumaCult alveolar organoid medium (*12*). The F medium + ROCK inhibitor was optimal for promoting epithelial cell attachment and enrichment of AT2 (HTII-280^+^) cells. In contrast, the serum-based medium yielded fewer AT2 cells, the B27-based medium supported good attachment but reduced AT2 cell retention (possibly due to differentiation), and the PneumaCult medium failed to support cell attachment and produced few AT2 cells (Figure 1D).

In 2D culture, these cell lines expressed HTII-280 and demonstrated the capacity to differentiate toward AT1 cells, with a subpopulation exhibiting transitional phenotypes marked by KRT8 and other known transitional cell markers (*13*). We performed single-cell RNA sequencing (scRNA-seq) on our healthy immortalized AT2 cell lines and integrated the data with published data (GSE132915) of primary AT2 cells as a control. This analysis demonstrated that 2D-cultured immortalized cells downregulated SFTPC (an AT2-specific marker), likely due to the lack of 3D cuboidal architecture, but retained other canonical markers, including LPCAT1 and ETV5. scRNA-seq identified distinct clusters corresponding to AT1-like cells and transitional states. Notably, a subset of cells expressed epithelial-mesenchymal transition (EMT) markers, indicating that these immortalized lines retain the potential to undergo mesenchymal transition, a feature relevant to fibrotic processes (Figure 1E). Immunofluorescence staining in 2D culture confirmed expression of the AT2 cell markers HTII-280 and LPCAT1, albeit with low levels of SFTPC. AT1 cell marker was detected by PDPN staining, while transitional cell markers (KRT8, KRT5, KRT17, and Claudin-4) showed robust expression (Figure 1F).

To maintain 3D cell structure, mimic the in vivo environment, and assess self-renewal and organoid formation capacity, we seeded immortalized cell lines in growth-factor– reduced Matrigel under feeder-free, serum-free 3D culture conditions (DMEM/F12 supplemented with B27, 20 ng/mL EGF, and 20 ng/mL FGF10) (*14*). Immortalized AT2 cell lines formed organoids efficiently, with higher colony-forming efficiency (CFE) in healthy lines than in IPF lines, confirming faithful mimicry of primary cell organoid-forming capacity and recapitulating the reduced regenerative capacity seen in primary IPF AT2 cells (Figure 2A). When we compared our conditions to Serum-Free, Feeder-Free (SFFF) medium which described by the Tata group (*15*), we found that our simplified medium (DMEM/F12 supplemented with B27, EGF, and FGF10) supported organoid formation with comparable CFE. This medium does not contain agents actively drive lineage specification pathways like WNT/β-catenin, TGF-β/BMP/SMAD or Hedgehog, making it more efficient for studying the effects of various inhibitors or treatments without interference from additional complex components (Figure 2B). Adding serum to the medium induced signs of EMT, whereas B27-supplemented medium strongly supported organoid formation.

**Figure 2:**
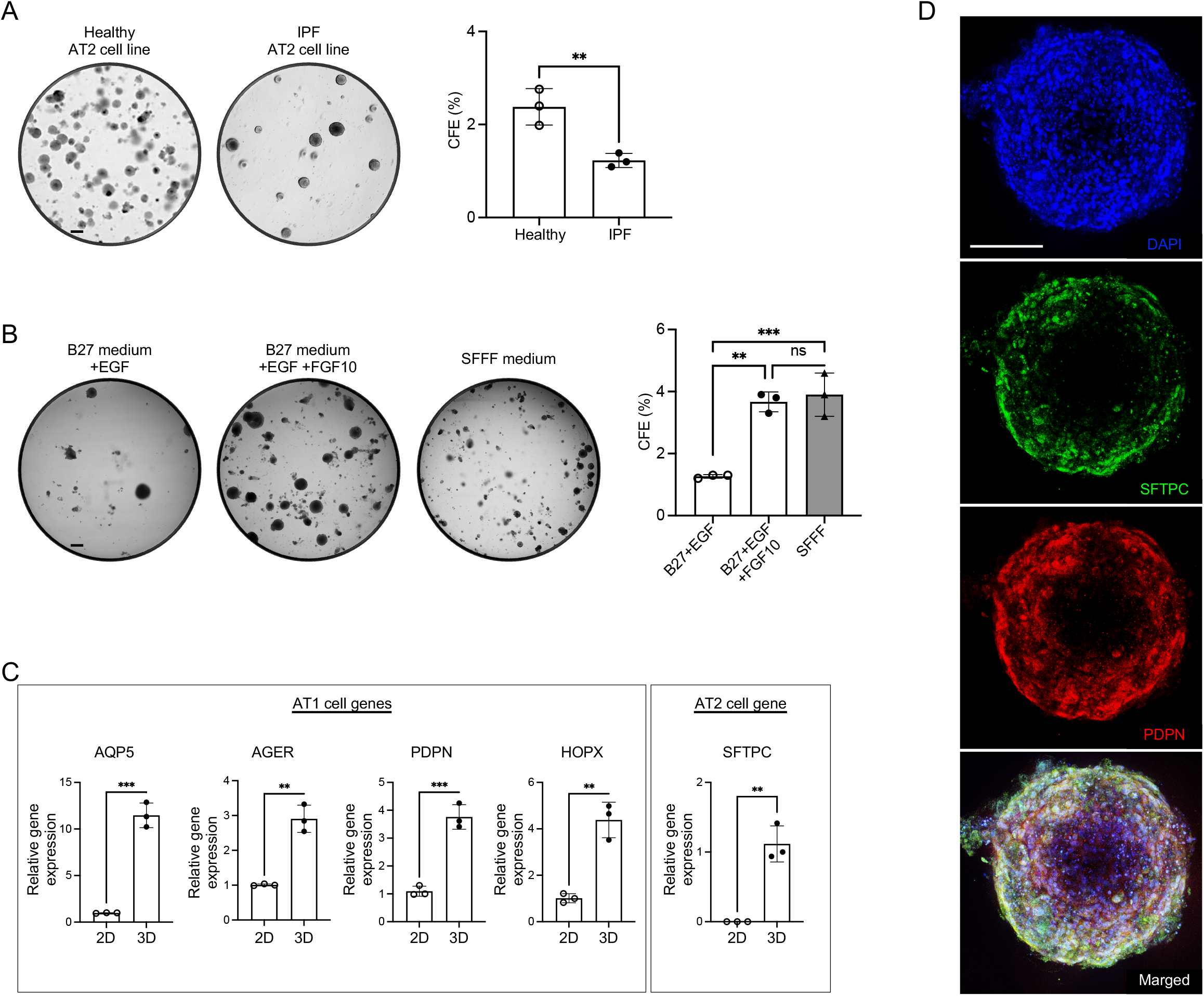
3D culture model of immortalized AT2 cell lines. (A) Representative images and colony-forming efficiency (CFE) of 3D organoid cultures (day 14) from healthy and IPF AT2 cell lines under feeder-free, serum-free 3D culture conditions (DMEM/F12 supplemented with B27, 20 ng/mL EGF, and 20 ng/mL FGF10). (n = 3 per group, **p < 0.01). Scale bars, 500 μm. (B) Representative images and CFE of 3D organoid cultures (day 14) from healthy AT2 cell lines under deferent feeder-free, serum-free 3D culture conditions: 1- B27 + EGF (DMEM/F12 supplemented with 1X B27, and 20 ng/mL EGF), 2- - B27 + EGF + FGF10 (DMEM/F12 supplemented with 1X B27, 20 ng/mL EGF, and 20 ng/mL FGF10), 3- SFFF medium (Advanced DMEM/F12, 1X Glutamax, 15mM HEPES, 10 μM SB431542, 3 μM CHIR99021, 1 μM BIRB796, 1X ITS, 1X B27, 1XN2, 50 ng/mL EGF, 1.25 mM N-Acetyl Cysteine, 10 μM Y27632, 10 ng/mL FGF10, 10 ng/mL hIL-1b). (n = 3 per group, **p < 0.01, ***p < 0.001). Scale bars, 500 μm. (C) Results from qPCR analysis of alveolar cell gene expression in healthy AT2 cell lines cultured under 2D or 3D conditions: *SFTPC* (AT2 marker) and *AQP5, AGER, PDPN*, and *HOPX* (AT1 markers). (n = 3 per group, **p < 0.01). (D) Immunofluorescence staining of whole organoids from healthy AT2 cell lines shows expression of AT2 cell marker SFTPC (green) and AT1 cell marker (red). Scale bars, 200 μm.

Analysis of AT1-specific genes revealed that the immortalized cell lines in 3D culture conditions exhibited enhanced differentiation toward AT1 cells. These cells also demonstrated increased functionality through upregulated SFTPC expression (Figure 2C). These findings were confirmed at the protein level using Immunofluorescence staining of organoids derived from the cell lines in the 3D model (Figure 2D).

In summary, we generated immortalized human AT2 cell lines from healthy and IPF lungs that recapitulate key features of alveolar epithelial biology in both 2D and 3D cultures, including self-renewal, differentiation, and transitional cell states. Optimized for serum-free, feeder-free organoid growth, these models provide an accessible platform to study distal lung regeneration, epithelial–mesenchymal interactions, and therapeutic responses in pulmonary disease. Their use may facilitate advances in drug discovery and personalized approaches for otherwise intractable lung disorders.

## Materials and Methods

### Generation and cultures of immortalized AT2 cell lines

Immortalized AT2 cell lines were generated using a modified protocol (*8*). Briefly, EpCAM^+^ cells were magnetically sorted from healthy and IPF lung tissues using CD326 (EpCAM) microbeads (Miltenyi Biotec, catalog 130-061-101). Sorted cells were resuspended in F medium + ROCK inhibitor (Y-27632) and allowed to adhere for 2 days. Adherent cells were then transduced with SV40 Large T antigen (SV40 LgT) lentivirus (customized from VectorBuilder) for immortalization, followed by selection with G418 (Sigma-Aldrich, Catalog G8168). Using FACS, immortalized AT2 (HTII-280^+^) cells were then sorted from the immortalized EpCAM^+^ population and cultured. To establish stable cell lines, the sorted immortalized AT2 cells were re-transduced with SV40 LgT. To investigate differences between various medium conditions, the immortalized AT2 cell lines were subsequently cultured under the following conditions: 1- F medium + ROCK inhibitor (3:1 DMEM/F12 to DMEM, 5% FBS, 0.4 μg/mL hydrocortisone, 5 μg/mL insulin, 8.4 ng/mL cholera toxin, 10 ng/mL EGF, and 10 μM Y-27632), 2- Serum-based medium (DMEM/F12, 10% FBS, 1X ITS and 10 μM Y-27632), 3- Serum-free (B27-based) medium (DMEM/F12 supplemented with 1X B27 and 10 ng/mL EGF + 10 μM Y-27632), 4- PneumaCult Alveolar Organoid medium.

### 3D organoid cultures of AT2 cells

Immortalized AT2 cell lines were cultured in a feeder-free 3D organoid condition as previously detailed (*14*). Briefly, cells were resuspended in growth factor-reduced Matrigel (Corning) and plated into a Matrigel-coated 24-well plate in domes (5 × 103 cells in 50 μl Matrigel/well). Cells were cultured under serum-free conditions consisting of DMEM/F12, supplemented with 1 × B-27 without vitamin A (Gibco, Catalog 12587010), 100 U/ml penicillin, 100 μg/ml streptomycin, 20 ng/ml human recombinant EGF (ThermoFisher, Catalog PHG0311), with or without 20 ng/ml human recombinant FGF10 (#100-26, Peprotech), or under SFFF medium (*15*). Colony formation efficiency (CFE%) was calculated as (number of organoids formed)/(number of cells seeded) × 100.

### Immunofluorescence staining

Cultured AT2 cell lines and organoids formed in 3D Matrigel were fixed with 4%paraformaldehyde (Sigma-Aldrich) in PBS. The 2D and 3D cultures were stained with primary antibodies: anti-HTII-280 (Terrace Biotech, #TB-27AHT2-280), anti-LPCAT1 (Proteintech, #16112-1-AP), anti-SFTPC (Proteintech, #10774-1-AP), anti-PDPN (DSHB, #8.1.1), anti-KRT5 (BioLegend, #905904), anti-KRT8 (DSHB, #TROMA-I), anti-KRT17 (Proteintech, #17516-1-AP), and anti-Claudin4 (Proteintech, #16195-1-AP). After washing, samples were incubated with secondary antibodies for 2 h at room temperature: Cy3 goat anti-rabbit IgG (Jackson Immunoresearch, #111-165-003), AF488 goat anti-mouse IgG/IgM (Invitrogen, #A-10680), AF594 goat anti-rabbit IgG (Invitrogen, #A-11012), AF488 goat anti-rat IgG (Invitrogen, #A-11006), or AF594 goat anti-hamster IgG (Invitrogen, #A-21113). Then samples were washed and stained with DAPI for 10 min. Fluorescence was imaged using a Nikon AX R NSPARC Confocal Microscope.

### Real-Time Quantitative PCR (qPCR)

RNA was isolated using a RNeasy Mini Kit (Qiagen) according to the manufacturer’s instructions. cDNA was prepared using a SuperScript VILO cDNA Synthesis Kit (Thermo Fisher Scientific). Real-time PCR was performed using Power SYBR Green PCR Master Mix (Applied Biosystems) on the ABI 7500 Fast Real-Time PCR System (Applied Biosystems). The specific primers are listed below: Human *SFTPC* forward ATCCCCAGTCTTGAGGCTCT and reverse CTTCCACTGACCCTGCTCAC. Human *PDPN* forward GGAAGGTGTCAGCTCTGCTC and reverse CGCCTTCCAAACCTGTAGTC. Human *AQP5* forward CGCTCAACAACAACACAACG, and reverse GAGTCAGTGGAGGCGAAGAT. Human *HOPX* forward TCAACAAGGTCGACAAGCAC, and reverse TCTGTGACGGATCTGCACTC. Human *AGER* forward GCCACTGGTGCTGAAGTGTA, and reverse TGGTCTCCTTTCCATTCCTG. Human *GAPDH* forward CCCATGTTCGTCATGGGTGT, and reverse TGGTCATGAGTCCTTCCACGATA. Relative gene expression was calculated using the ΔΔCt method and normalization to two housekeeping genes.

### Statistics

Statistical analysis was performed using the GraphPad Prism software, using Student’s t-test and ANOVA. Results were considered statistically significant at **P < 0.01, and ***P < 0.001. The number of independent biological replicates is reported as n.

## Acknowledgements

The authors thank the members of the Jiang-Liang Laboratory and the lung institute for helpful discussion.

Supported by National Institutes of Health (NIH) grants R01 HL172990 (D.J.) and R01 AG078655 (J.L.).

## Author Contribution

A.R. performed experiments, analyzed and visualized data, and wrote the manuscript. Y.Q. and X.Z. performed experiments. J.L. and D.J. supervised the project and wrote the manuscript. All authors contributed to the paper, and read and approved the submitted version.

## Data availability

Single-cell RNA sequencing data were deposited and are available by contacting the corresponding author.

